# A Data-Driven Correction Framework for Axial- and Radial-Position-Dependent Intensity Attenuation in Volumetric Fluorescence Microscopy

**DOI:** 10.64898/2026.05.24.727559

**Authors:** Saya Ichihara, Shotaro Akaho, Satoshi Kuriki, Kohei Otomo, Tomomi Nemoto, Akatsuki Kimura

## Abstract

Accurate quantification of fluorescence signals in three-dimensional (3D) microscopy is often hindered by axial- and radial-position-dependent attenuation, limiting reliable measurements in live biological specimens. Here, we present a data-driven statistical correction model that compensates for signal loss arising from axial- and radial-position in 3D time-lapse imaging of *Caenorhabditis elegans* embryos. Our framework incorporates axial position (imaging depth, *z*), radial position (distance from the center of the field of view, *r*), together with cell cycle progression, to recover cell-specific fluorescence intensities independent of axial- and radial-positions. By leveraging repeated observations of biologically comparable states, the model infers attenuation directly from the data without requiring external calibration. Notably, the sign of the inferred radial-position-dependence in biological specimens was opposite to that observed in homogeneous fluorescent reference samples, underscoring the value of specimen-specific, data-driven correction. Validation using histone-tagged fluorescent proteins demonstrated that the method effectively removes geometric bias in nuclear fluorescence signals, enabling consistent quantification across cells and embryos. This approach provides a robust and generalizable solution for correcting intensity attenuation in volumetric microscopy datasets, thereby enabling more accurate and reproducible quantitative analyses in live imaging studies.

**Author summary:** Modern microscopy lets us watch living cells and embryos in three dimensions, but measuring brightness accurately is harder than it seems. Signals often become weaker not only when molecules are less abundant, but also when they lie deeper in the specimen or farther from the center of the image. This makes it difficult to tell whether differences in brightness reflect biology or simply the position of a cell within the microscope field. Existing correction methods typically rely on separately acquired reference measurement samples or on image-level statistical patterns. In this study, we took a different approach. Using the highly reproducible development of nematode (*C. elegans*) embryos, we compared cells that should be biologically equivalent across multiple embryos and used those repeated observations to estimate imaging bias directly from the biological images themselves. Our method corrects for both axial-dependent and radial attenuation simultaneously within a unified statistical framework, requiring no such reference data. Beyond simply improving consistency, we uncovered an unexpected result: the radial bias inferred from real embryos was opposite in sign to what calibration samples would predict. This underscores the need for specimen-specific, data-driven correction. Our framework should help make live imaging more quantitatively accurate for studying dynamic biological processes in complex three-dimensional specimens.

## Introduction

Fluorescence microscopy is a cornerstone of modern cell and developmental biology, enabling the direct measurement of molecular and cellular dynamics in living systems [1]. In particular, confocal microscopy based on single-photon excitation has become one of the most widely used tools, allowing researchers to visualize the spatial distribution of molecules in cells and tissues with relative ease.

Fluorescence intensity is, in principle, proportional to the number of fluorescent molecules, suggesting that microscopy could provide a direct quantitative readout of molecular abundance. Such quantitative information would enable rigorous comparisons across cells, developmental stages, and experimental conditions. However, in three-dimensional (3D) imaging, this assumption often breaks down due to non-uniform attenuation of fluorescence intensity caused by scattering, absorption, and optical aberrations within thick specimens. As a result, fluorescence intensity depends not only on molecular abundance but also on the axial position (*z*; imaging depth along the optical axis) and the radial position (*r*; distance from the center of the image, which coincides with the optical axis). In practice, signals become progressively dimmer with increasing *z*, even when the underlying molecular amount is identical [2]. These imaging-dependent effects introduce systematic bias into quantitative measurements and complicate the distinction between biological variability and imaging-induced artifacts.

Several classes of approaches have been explored to overcome these limitations. A first direction is to improve imaging hardware or acquisition settings to reduce signal attenuation. Two-photon microscopy confines excitation to the focal volume and reduces scattering due to the use of longer excitation wavelengths, making it advantageous for imaging thick specimens [3]. Light-sheet microscopy enables optical sectioning with reduced photodamage and a reduced dependence of signal on axial position (*z*) [4,5]. A simpler and more widely applicable approach is to progressively increase laser power with imaging depth within a conventional confocal microscope, partially compensating for signal loss along the z-axis [6]. However, none of these strategies fully eliminates position-dependent attenuation: two-photon excitation remains limited by scattering of ballistic photons [2,7], light-sheet microscopy remains susceptible to absorption and scattering within the specimen [4], and empirical laser power compensation leaves residual intensity gradients [6]. Furthermore, the hardware-based approaches in particular require specialized or expensive equipment that is not universally accessible.

A second direction is to estimate and correct for signal attenuation computationally. One class of approaches relies on physical modeling of optical effects, incorporating light attenuation, point spread function (PSF) variations, or scattering properties into correction algorithms [8–10]. These methods are theoretically grounded and can be effective when the optical properties of the sample are well characterized. However, deriving accurate optical parameters for heterogeneous biological specimens is generally impractical, limiting the applicability of such approaches in live imaging contexts.

A complementary class of approaches instead infers the attenuation function empirically from the data itself, without explicit physical modeling. Early implementations estimated the *z*-dependent decay by fitting a parametric function — typically exponential — to the mean fluorescence intensity computed across all pixels in each z-plane [11]. While this avoids the need for optical parameters, it rests on the assumption that fluorophore density is approximately uniform along the axial direction, which is rarely satisfied in structured biological specimens where the spatial distribution of fluorescent molecules is inherently heterogeneous.

A more refined variant of this empirical strategy was developed in the context of quantitative cell biological analysis of *Caenorhabditis elegans* embryos [6]. Rather than relying on the imaged specimen itself, this approach uses a panel of 11 reference strains expressing fluorescent proteins at ubiquitous and nominally constant levels, fitting the relationship between fluorescence intensity and z-position to derive attenuation correction functions. Although this substantially improved quantitative consistency across embryos, the method requires extensive prior preparation of dedicated reference strains and assumes that the selected proteins are expressed at truly constant levels across all cells — an assumption that is difficult to verify independently. Furthermore, it addresses only axial (*z*) attenuation and does not account for radial (*r*) position-dependent variation.

To address these limitations, we developed a data-driven statistical framework that estimates and corrects for both axial- (*z*) and radial- (*r*) position-dependent fluorescence attenuation directly from replicated observations of the biological specimen of interest, without requiring dedicated reference strains or prior optical calibration. Unlike previous approaches, which typically estimate correction factors from a single sample, a predefined optical model, or a panel of external reference strains, our method explicitly leverages repeated imaging of the same biological system across multiple specimens acquired at different spatial positions. We assume that fluorescence intensity should be equivalent across cells sharing the same identity and cell-cycle stage, and use this constraint – applied across multiple embryos imaged at varying axial and radial positions – to infer the dependence of observed signal on imaging geometry directly from the data. By integrating information across replicated measurements, the framework separates cell-intrinsic fluorescence from imaging-induced variation, without assuming a specific functional form for the attenuation: candidate models are evaluated and selected using the Akaike Information Criterion (AIC), allowing the structure of the correction to be determined empirically. We applied this approach to *C. elegans* embryos, where the high reproducibility of embryogenesis and well-defined cell identities make it possible to identify biologically equivalent cells across specimens. Using histone-tagged fluorescent proteins as a benchmark, we demonstrate that the method successfully recovers cell-intrinsic fluorescence intensity independent of imaging geometry. Because the framework does not depend on specific imaging modalities or prior optical calibration, it can in principle be extended to other biological systems where equivalent cellular states can be defined across replicated observations.

## Results

### Evaluation of axial-position-dependent intensity attenuation and nuclear morphology in one- and two-photon spinning-disk confocal microscopy

We first evaluated whether improvements in imaging hardware can mitigate geometric bias in fluorescence quantification. We extended a conventional spinning-disk confocal microscope to a two-photon excitation configuration [12–14]. In this system, excitation is achieved either by one-photon (488/561 nm) or two-photon (920/1064 nm) illumination, while the downstream optical path, including the spinning-disk confocal unit (Yokogawa CSU-X1-base), is identical. Thus, any differences in imaging performance can be attributed primarily to the excitation modality.

As a test system, we analyzed *C. elegans* embryos expressing histone-fused fluorescent proteins (GFP::H2B or mCherry::H2B) (Fig. 1A–D). Histone signals provide a suitable benchmark for quantitative evaluation because they are localized within nuclei, allowing clear spatial definition of measurement regions, and are expected to exhibit relatively small variability in fluorescence intensity across cells within the same developmental stage. This makes it possible to assess whether observed intensity variation reflects biological differences or imaging-induced artifacts. Nuclear segmentation for quantitative analysis was performed using Imaris (see Methods for details) (Fig. 1A’–D’). The fluorescence intensity quantified in this study corresponds to the mean intensity within each segmented nucleus, which reflects the nuclear concentration of histone-associated fluorescence rather than total nuclear fluorescence. This pipeline enabled quantitative comparison of nuclear fluorescence intensity and morphology across imaging conditions.

**Figure 1:**
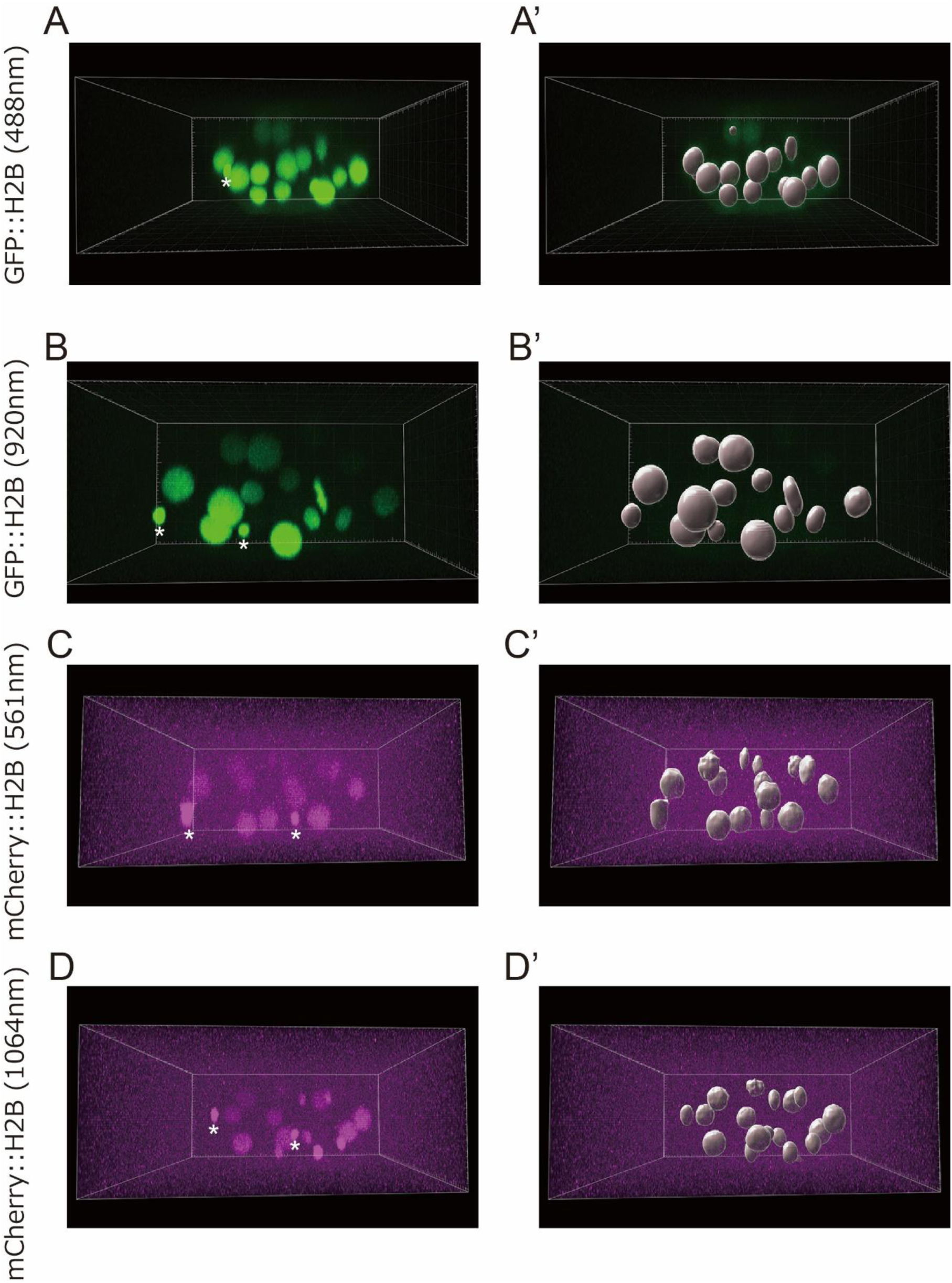
Representative three-dimensional images of a *C. elegans* embryo expressing histone proteins at the 14-cell stage. Images were reconstructed in three dimensions using Imaris and are shown from the lateral view along the short axis of the embryo (y-axis). The bottom side of each image corresponds to the side closer to the objective lens. (A) Embryo expressing GFP::H2B imaged at 488 nm (one-photon excitation). (B) Embryo expressing GFP::H2B imaged at 920 nm (two-photon excitation). (C) Embryo expressing mCherry::H2B imaged at 561 nm (one-photon excitation). (D) Embryo expressing mCherry::H2B imaged at 1064 nm (two-photon excitation). Asterisks indicate polar body. (A’-D’) Corresponding segmented images generated in Imaris. Nuclear regions were segmented in three dimensions, and each nucleus was assigned a cell identity.

We first examined axial-position(*z*)-dependent changes in fluorescence intensity. In both imaging modalities, a prominent decrease in fluorescence intensity along the *z*-axis was observed (Fig. 1A–D). Quantitative analysis confirmed that fluorescence intensity systematically decreased with *z* in both modalities (Fig. 2A–D), indicating that axial-position-dependent attenuation persists even under two-photon excitation.

**Figure 2:**
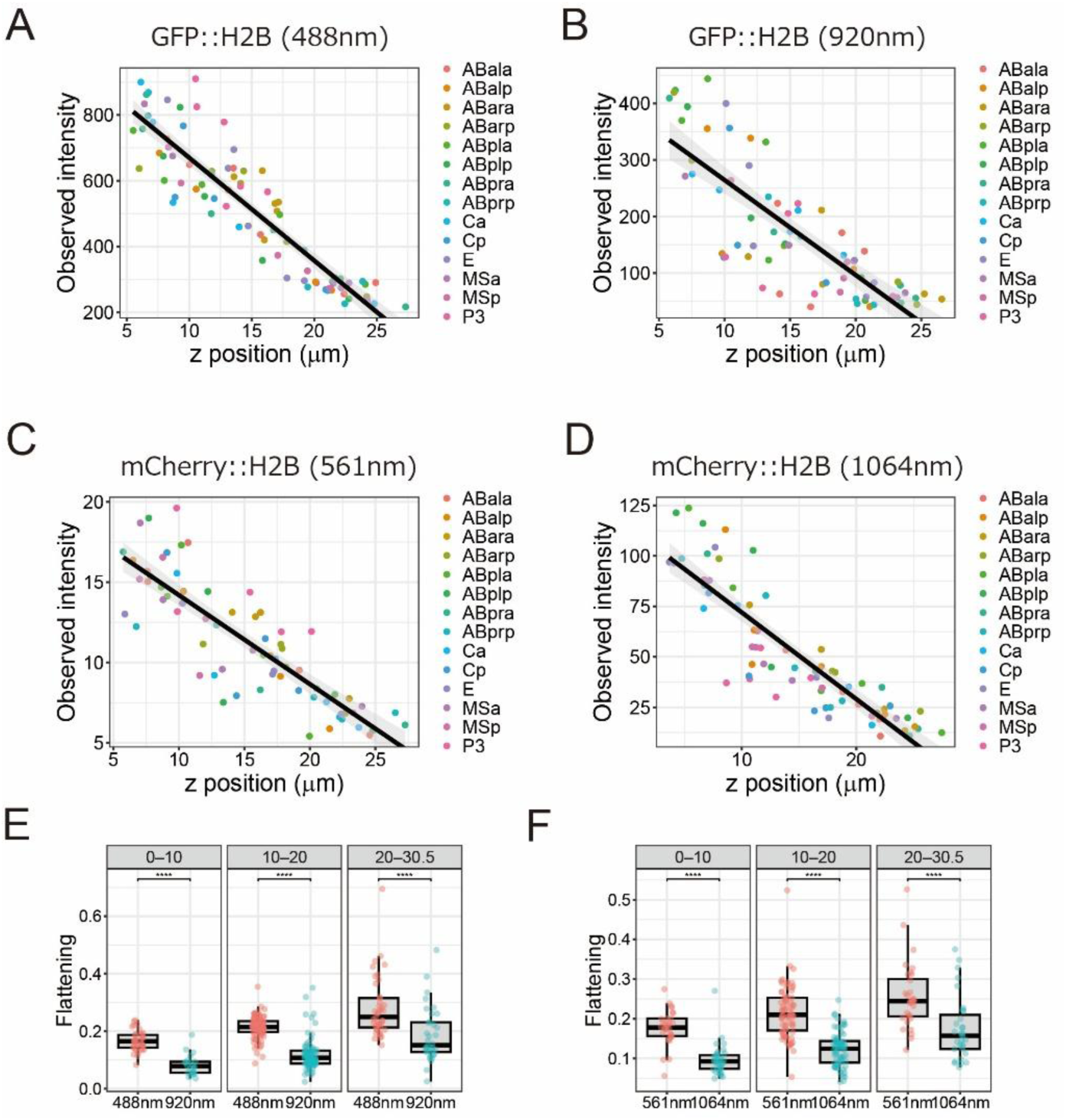
Axial-position-dependence of signal intensity and nuclear shape. (A–D) Axial-position-dependence of nuclear fluorescence intensity. Nuclear mean intensity at the 14-cell stage (*T* = 0.5, corresponding to the midpoint of the cell cycle) was plotted against the z-position of each nucleus. (A) GFP::H2B imaged at 488 nm (slope = −0.067, *R*^2^ = 0.86). (B) GFP::H2B imaged at 920 nm (slope = −0.105, *R*^2^ = 0.68). (C) mCherry::H2B imaged at 561 nm (slope = −0.052, *R*^2^ = 0.76). (D) mCherry::H2B imaged at 1064 nm (slope = −0.093, *R*^2^ = 0.83). Black lines indicate linear regression fits, and gray shaded areas represent confidence intervals. (E) Comparison of nuclear flattening between one-photon and two-photon imaging for GFP::H2B at the 14-cell stage (*T* = 0.5). Flattening was defined as 1 – *a*/*b*, where *a* denotes the mean of the two minor-axis radii projected onto the x-y plane, and *b* denotes the major-axis radius projected onto the z-axis. Data were grouped into three bins based on *z*-position, and flattening values were compared within each bin. Pink and light blue indicate one-photon and two-photon imaging, respectively. Statistical significance was assessed using Wilcoxon rank-sum tests with Holm correction for multiple comparisons (\**p* < 0.05; \*\**p* < 0.01; \*\*\**p* < 0.001; \*\*\*\**p* < 0.0001). (F) Same analysis as in (E) for mCherry::H2B at the 14-cell stage (*T* = 0.5). Data were collected from six embryos for GFP::H2B imaged at 488 nm and from five embryos for each of the other imaging conditions.

We next assessed nuclear morphology as an indicator of imaging fidelity. Visual inspection of nuclei after segmentation in Imaris revealed differences between the two imaging modalities. Nuclei imaged with two-photon microscopy appeared relatively spherical, whereas those imaged with one-photon microscopy appeared elongated along the *z*-axis (Fig. 1A’–D’). To quantify this effect, we calculated nuclear flattening, defined as the relative reduction of nuclear diameter along the *z*-axis compared to the lateral diameter (with 0 indicating isotropic shape). Nuclei imaged with two-photon microscopy consistently showed lower flattening across all axial positions, indicating better preservation of their native shape (Fig. 2E,F and Sup. Fig. S1). This improvement is consistent with previous optical characterizations showing that, in similar spinning-disk configurations, two-photon excitation yields a narrower axial point spread function than one-photon excitation [12]. These experiments confirmed that two-photon microscopy is better suited for three-dimensional imaging in terms of segmentation accuracy.

Although two-photon microscopy improves the geometric fidelity of nuclear morphology, it does not resolve the problem of intensity attenuation: fluorescence signals still decrease systematically with increasing *z* position (Fig. 2A–D). These results indicate that hardware-based improvements alone are insufficient to eliminate axial-position-dependent bias in fluorescence intensity measurements.

In addition, fluorescence intensity should also be considered as a function of radial-position (*r*), even within the same focal plane. We observed that, when a homogeneous fluorescent plate was imaged, fluorescence intensity decreased toward the periphery of the field of view (Fig. S2A–D). This *r* bias arises from the optical configuration particularly in our two-photon system, in which the beam expander was adjusted to concentrate excitation power near the optical axis. Taken together, these observations demonstrate that fluorescence measurements in volumetric imaging are influenced by both axial- (*z*) and radial- (*r*) positions. Therefore, accurate quantitative analysis requires a correction framework that simultaneously accounts for these geometric dependencies.

### Development of a statistical correction model for geometric attenuation

To correct fluorescence intensity biases arising from axial- (*z*) and radial- (*r*) positions, we developed a data-driven statistical framework that estimates geometric attenuation directly from volumetric imaging data. The key idea of our approach is to leverage repeated imaging of multiple samples to identify spatially dependent biases. Specifically, we assume that fluorescence intensity should be comparable across equivalent biological states and use measurements of such equivalent points acquired at different spatial positions to infer attenuation as a function of imaging geometry. In this study, we applied this framework to *C. elegans* embryos, where cell identities are well defined. This allows us to treat nuclei belonging to the same cell identity at comparative cell cycle timing as having similar underlying fluorescence intensity between different embryos.

Based on this idea, we constructed a statistical model that captures the dependence of observed fluorescence intensity on *z* and *r*, while separating these effects from biological variation (see Methods for details). For parameter estimation, we modeled the mean fluorescence intensity, *I*_i_, as:

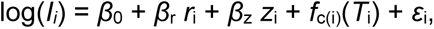

where *r*_i_ denotes the radial position, *z*_i_ denotes the normalized axial position, and *T*_i_ represents normalized time. The function *f*_c(i)_(*T*_i_) captures cell-specific temporal variation, modeled using natural cubic splines, and *ε*_i_ represents residual variation. In practice, we also considered extended models incorporating nonlinear *z* dependence and varying levels of temporal flexibility, as described below. This formulation enables the separation of geometric attenuation from biological variability using observational data alone.

### Model selection based on AIC

To determine the appropriate model structure, we compared a set of candidate models incorporating different combinations of *z*, *r*, and temporal (*T*) effects. Specifically, we evaluated models with or without *r* dependence, and with *z* dependence modeled as no effect, a linear term, or a quadratic term. Temporal variation was modeled using natural cubic splines with different degrees of flexibility. All candidate models were fitted to the log-transformed fluorescence intensity, and model performance was evaluated using the Akaike Information Criterion (AIC) [15]. The model with the lowest AIC was selected as the best representation of the data (Sup. Tables S1–S4).

Model selection consistently supported the inclusion of *z* dependence across all datasets, with the best-fitting models incorporating both linear and quadratic terms (Fig. 3A–D; Sup. Table S1–S4). Similarly, *r* dependence was also supported in all datasets. Temporal effects were selected equally between the two-knot model and the three-knot model. This supports the value of varying the number of knots for evaluation. This model selection procedure allows us to determine, in a data-driven manner, whether such dependencies are required and how they should be parametrized, without imposing these assumptions a priori.

**Figure 3:**
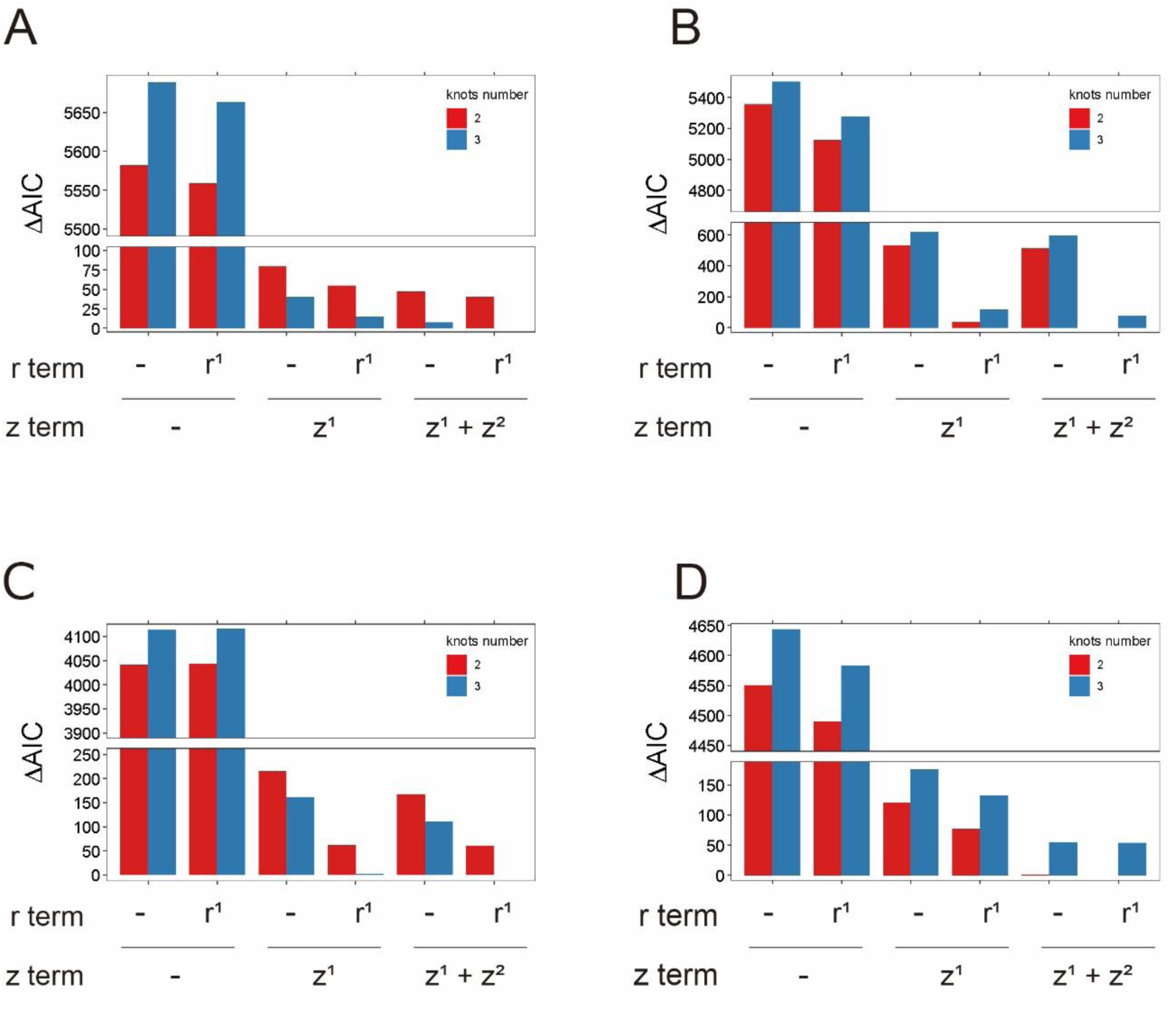
Comparison of candidate models based on AIC within each dataset. The difference in Akaike Information Criterion (ΔAIC) relative to the best-fitting model is shown for all 12 candidate models. Low ΔAIC values indicate better model fit. Model conditions represent combinations of radial dependence (r0: no radial term; r1: with radial term), axial dependence (z0: no *z* term; z1: linear *z* term; z12: linear and quadratic *z* terms), and the number of spline knots (2 or 3) used to model temporal effects. (A) GFP::H2B imaged at 488 nm. (B) GFP::H2B imaged at 920 nm. (C) mCherry::H2B imaged at 561 nm. (D) mCherry::H2B imaged at 1064 nm.

### Unexpected behavior of radial dependence in biological samples

We next examined the estimated coefficients for *r* dependence (*β*_r_) (Table 1). In all datasets, *β*_r_ was positive, indicating that fluorescence intensity increased with *r*. This behavior contrasts with calibration measurements using a homogeneous fluorescent plate (Sup. Fig. S2), where fluorescence intensity decreased with *r*, particularly in two-photon imaging. The discrepancy indicates that *r* dependence observed in biological samples cannot be explained solely by optical properties measured in homogeneous samples. These results suggest that attenuation should be inferred directly from biological data, supporting the utility of the present data-driven approach.

**Table 1.**
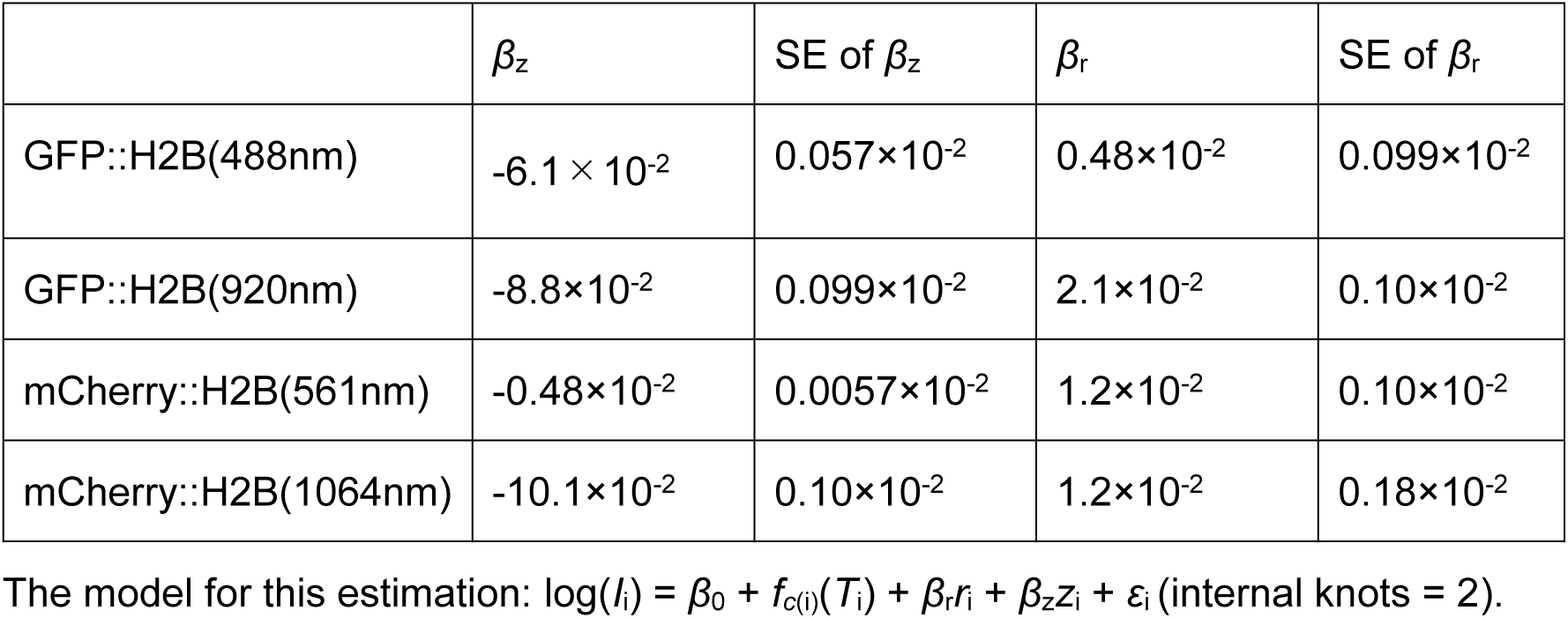
The estimated parameters for *z* and *r*-dependency and their standard errors (SEs)

| | $\beta_z$ | SE of $\beta_z$ | $\beta_r$ | SE of $\beta_r$ |
| --- | --- | --- | --- | --- |
| GFP::H2B(488nm) | $-6.1 \times 10^{-2}$ | $0.057 \times 10^{-2}$ | $0.48 \times 10^{-2}$ | $0.099 \times 10^{-2}$ |
| GFP::H2B(920nm) | $-8.8 \times 10^{-2}$ | $0.099 \times 10^{-2}$ | $2.1 \times 10^{-2}$ | $0.10 \times 10^{-2}$ |
| mCherry::H2B(561nm) | $-0.48 \times 10^{-2}$ | $0.0057 \times 10^{-2}$ | $1.2 \times 10^{-2}$ | $0.10 \times 10^{-2}$ |
| mCherry::H2B(1064nm) | $-10.1 \times 10^{-2}$ | $0.10 \times 10^{-2}$ | $1.2 \times 10^{-2}$ | $0.18 \times 10^{-2}$ |

The model for this estimation: $\log(I_i) = \beta_0 + f_{c(i)}(T_i) + \beta_r r_i + \beta_z z_i + \varepsilon_i$ (internal knots = 2).

### Correction reduces spatial dependence of fluorescence intensity

Using the best-fitting model selected as described above, we applied the correction framework to histone fluorescence data. To evaluate the effectiveness of the correction, we analyzed residuals of the fitted model as a function of imaging *z* and *r* (Fig. 4 and 5; *p* > 0.8 for testing slope = 0 in all cases). The residuals, aggregated across all time points, exhibited no detectable trends with respect to either *z* or *r*, indicating that the dominant spatial biases had been effectively removed (Fig. 4A–D and Fig. 5A–D). In contrast, models that did not include axial-position-dependent terms showed clear residual structure along the *z*-axes (Fig. 4A’–D’; the slope was significantly different from zero (*p* < 0.05) in all cases), demonstrating that *z* dependence is essential to account for geometric attenuation. Exclusion of *r* terms resulted in more subtle residual patterns (Fig. 5A’–D’; *p* < 0.05 in Fig. 5B’, 5C’ and 5D’; *p* = 0.17 in 5A’), consistent with the weaker but non-negligible contribution of *r* (Fig. 3A–D). The absence of structure in the residuals provides a critical validation of the model, demonstrating that the correction successfully captures the major sources of spatial bias. Partial *R*^2^ analysis quantitatively supported the observations that depth (*z*) contributes dominantly to variance compared to radial position (*r*) (Sup. Table S5).

**Figure 4:**
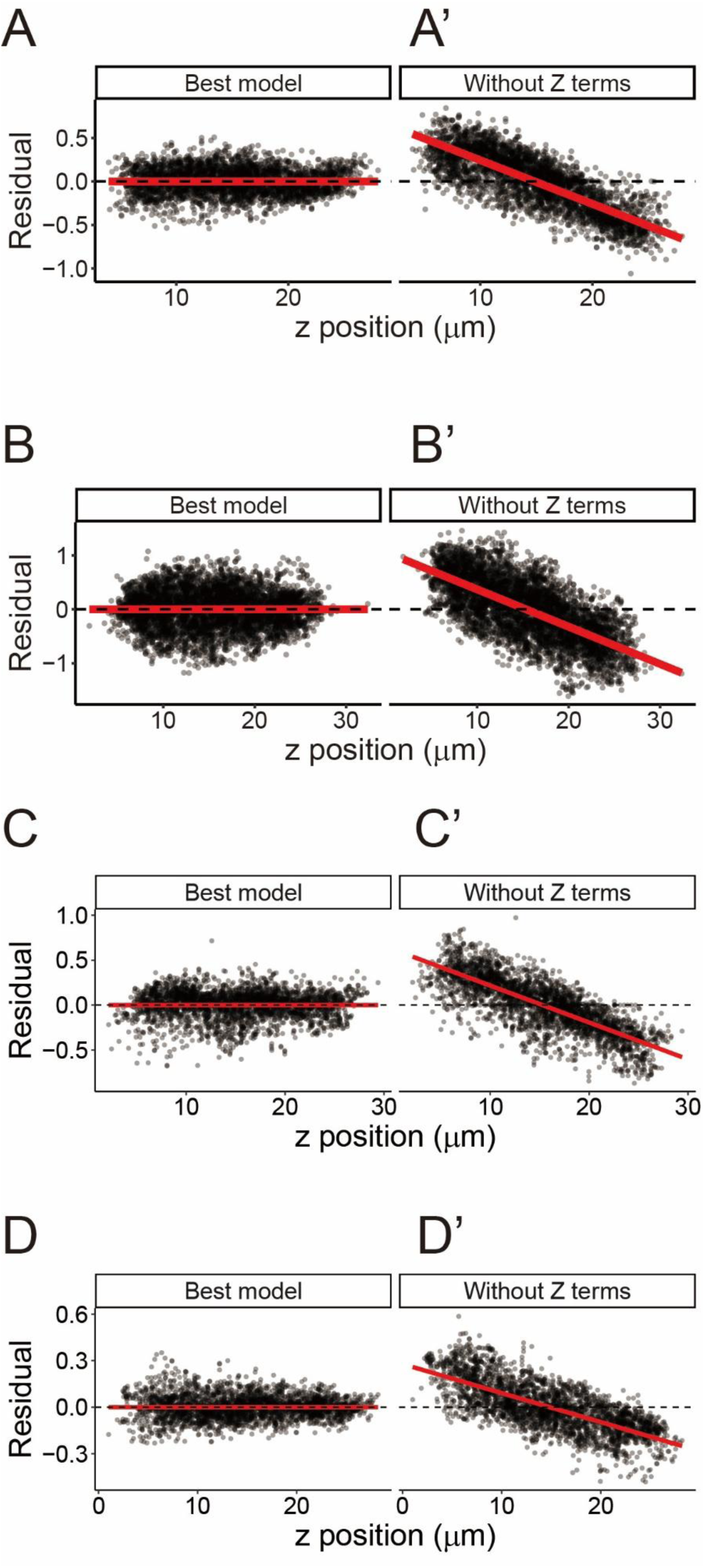
Residual dependence on axial position. (A-D) Residuals from the best-fitting model plotted against axial position (*z*). (A) GFP::H2B imaged at 488 nm (slope = −8.3 × 10⁻¹⁸, *R*^2^ = 0). (B) GFP::H2B imaged at 920 nm (slope = −4.3 × 10⁻¹⁸, *R*^2^ = 0). (C) mCherry::H2B imaged at 561 nm (slope = −1.0 × 10⁻^17^, *R*^2^ = 0). (D) mCherry::H2B imaged at 1064 nm (slope = −6.0 × 10⁻¹⁸, *R*^2^ = 0). (A′–D′) Residuals from models lacking axial-position-dependent terms, plotted against axial position (*z*). (A′) GFP::H2B imaged at 488 nm (slope = −0.050, *R*^2^ = 0.65). (B′) GFP::H2B imaged at 920 nm (slope = −0.069, *R*^2^ = 0.49). (C′) mCherry::H2B imaged at 561 nm (slope = −0.041, *R*^2^ = 0.2). (D′) mCherry::H2B imaged at 1064 nm (slope = −0.019, *R*^2^ = 0.53). Each point represents the residual for a single datapoint, and red lines indicate linear regression fits.

**Figure 5:**
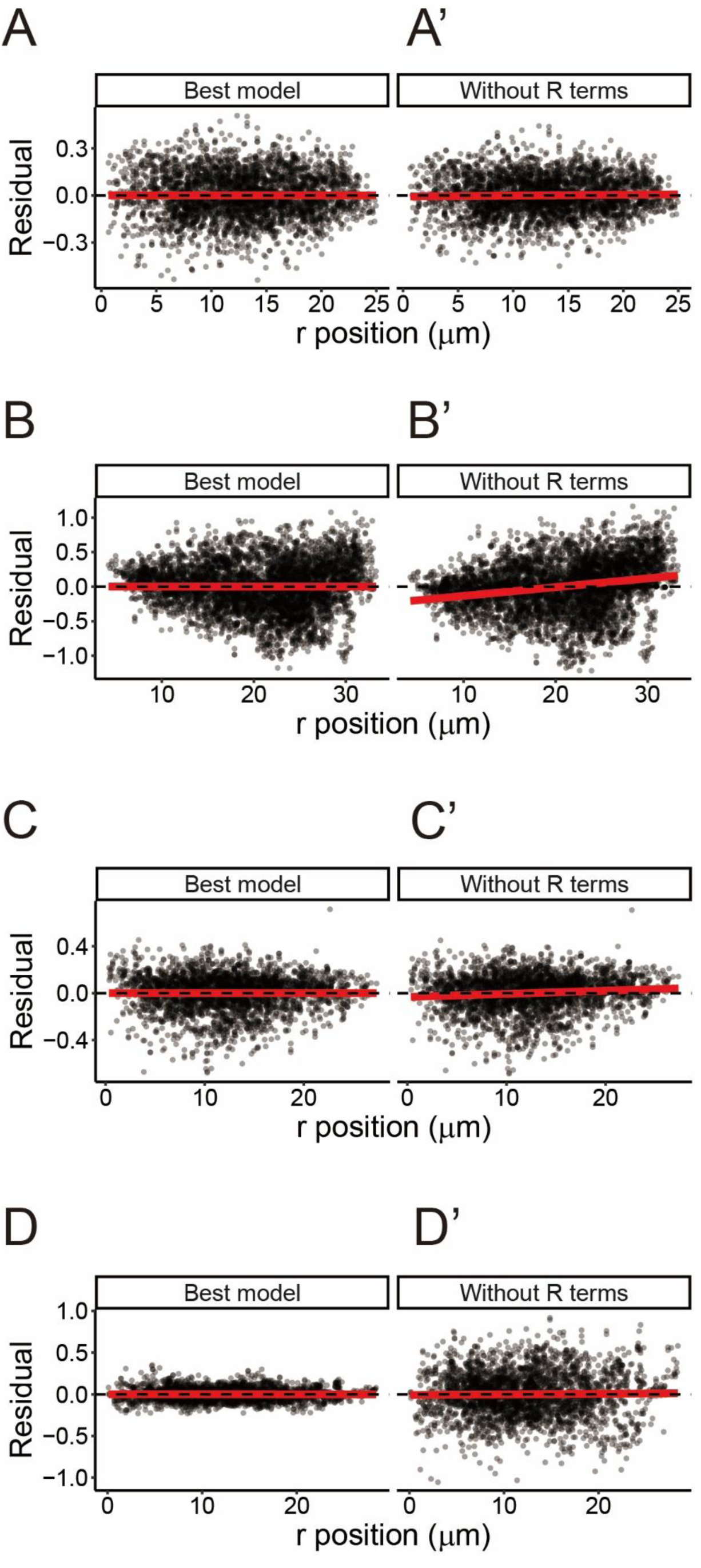
Residual dependence on radial position. (A–D) Residuals from the best-fitting model plotted against radial distance (*r*). (A) GFP::H2B imaged at 488 nm (slope = -4.5 × 10⁻^18^, *R*^2^ = 0). (B) GFP::H2B imaged at 920 nm (slope = -2.3 × 10⁻¹^9^, *R*^2^ = 0). (C) mCherry::H2B imaged at 561 nm (slope = - 2.4 × 10⁻¹^8^, *R*^2^ = 0). (D) mCherry::H2B imaged at 1064 nm (slope = -6 × 10^-5^, *R*^2^ = 0.2 × 10^-4^). (A′–D′) Residuals from models lacking radial term, plotted against radial distance. (A′) GFP::H2B imaged at 488 nm (slope = 5.3 × 10⁻⁴, *R*^2^ = 5.4 × 10^-4^). (B′) GFP::H2B imaged at 920 nm (slope = 0.013, *R*^2^ = 0.053). (C′) mCherry::H2B imaged at 561 nm (slope = 2.7 × 10⁻^3^, *R*^2^ = 0.009). (D’) mCherry::H2B imaged at 1064 nm (slope = 6.9 × 10^-4^, *R*^2^ = 0.0002). Each point represents the residual for a single datapoint, and red lines indicate linear regression fits.

### Correction reveals cell-cycle–dependent dynamics

We next examined the temporal dynamics of histone fluorescence intensity within a single cell cycle as revealed after correction. Here we focused on the 8-cell stage (Fig. 6) as a representative example, as comparable patterns were consistently observed across developmental stages.

**Figure 6:**
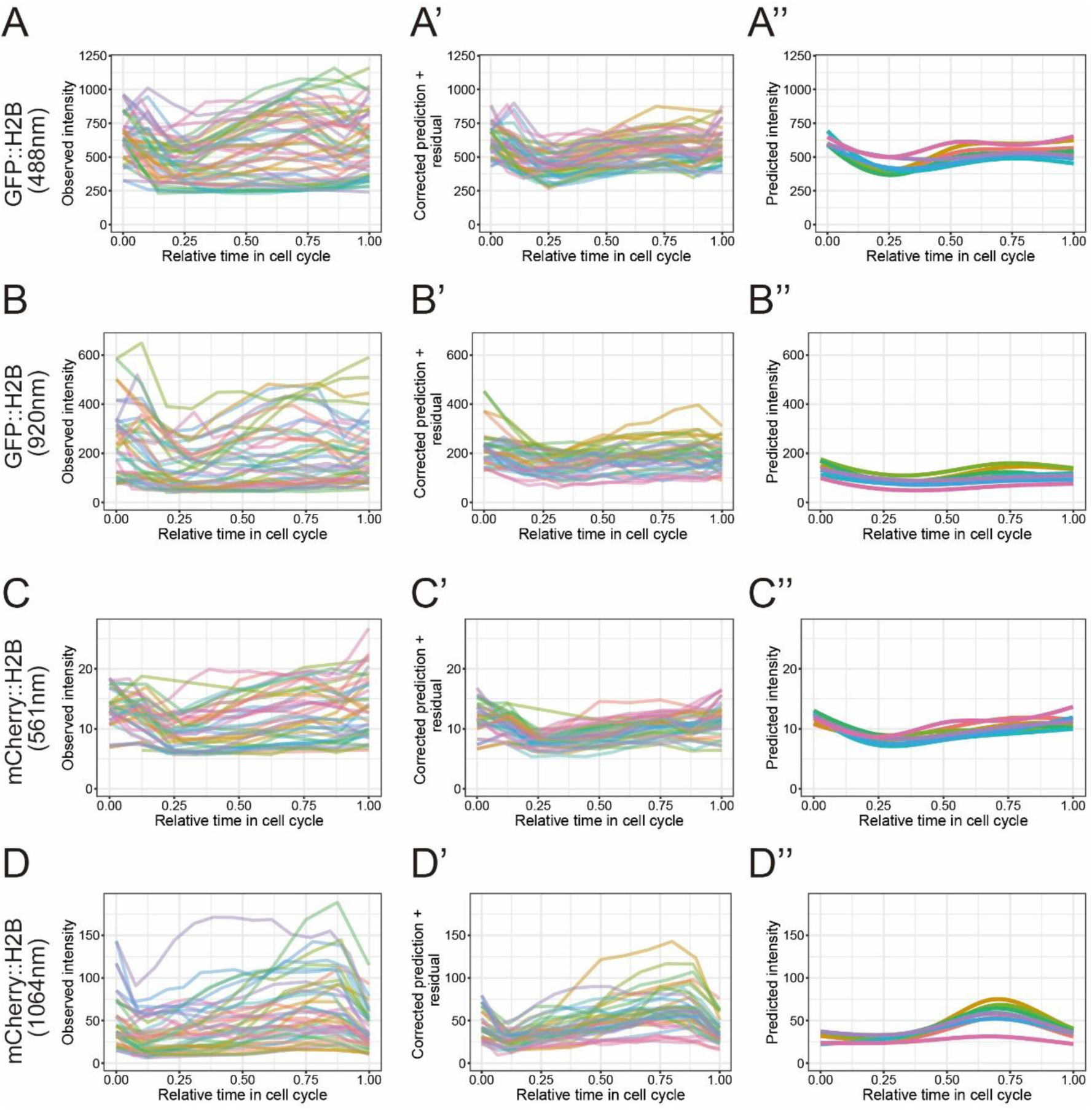
Cell cycle-dependent fluorescence dynamics at the 8-cell stage. (A–D) Observed fluorescence intensities plotted as a function of normalized time (*T*). (A) GFP::H2B imaged at 488 nm. (B) GFP::H2B imaged at 920 nm. (C) mCherry::H2B imaged at 561 nm. (D) mCherry::H2B imaged at 1064 nm. (A′–D′) Corrected fluorescence intensities including residual variation (predicted intensity + residual), plotted as a function of normalized time. These values correspond to intensities normalized to a common reference position (*r* = 0, z = embryo center). (A″–D″) Model-derived fluorescence intensities excluding residual variation, representing the estimated underlying cell-type-specific dynamics. In panels (A–D) and (A′–D′), each line represents an individual embryo, whereas in panels (A″–D″), each line represents the mean trajectory for a given cell type. Each color represents a distinct cell: ABal (salmon pink), ABar (ochre), ABpl (yellow-green), ABpr (green), C (light blue), E (blue), MS (purple), and P3 (pink).

We first compared raw fluorescence intensities (Fig. 6A–D) with corrected intensities that retain residual variability (Fig. 6A′–D′). In the raw data, substantial variability was observed across embryos even for the same cell type (Fig. 6A–D). After correction, the overall spread of the trajectories became narrower, extreme deviations were largely reduced, and the temporal profiles became more similar across cells (Fig. 6A’–D’). Furthermore, no trend was observed when the residuals were plotted against time (Sup. Fig. S3). These changes indicate that a substantial portion of the observed variability originates from imaging-dependent bias rather than biological differences.

We then examined the model-derived cell-specific intensity trajectories, in which residual noise was removed to estimate the underlying fluorescence dynamics (Fig. 6A”–D”). These trajectories revealed a consistent U-shaped temporal profile across all datasets, characterized by an initial decrease followed by a gradual increase. The initial decrease in mean fluorescence intensity (*T* < ∼0.25) coincides with an increase in nuclear volume and is consistent with dilution of histone concentration due to nuclear expansion after mitosis (Sup. Fig. S4A, B). In contrast, the subsequent increase in mean intensity despite continued nuclear growth indicates an increase in total histone content within the nucleus. This increase is likely attributable to processes such as nuclear import of histones, DNA replication and chromatin condensation. Thus, the corrected intensities recover biologically interpretable dynamics that are obscured in the raw data.

### Correction reveals stage-invariant intensity and reduces variability across embryos

We next examined how the correction affects variability across cells at the same developmental stages. Before correction, observed fluorescence intensities showed substantial variability across cells even within the same stage (2-, 4-, 8-, and 14-cell stages), reflecting the combined effects of biological variation and imaging-induced bias (Fig. 7A–D). In contrast, corrected intensities exhibited markedly reduced variability and converged to nearly constant values within each stage (Fig. 7A’–D’). Importantly, in our model, fluorescence intensities were not constrained to be identical across different cells at the same developmental stage. The emergence of stage-wise consistency is therefore not a trivial consequence of the model assumptions but rather indicates that geometric biases have been effectively removed and that the remaining variation reflects underlying biological structure.

**Figure 7:**
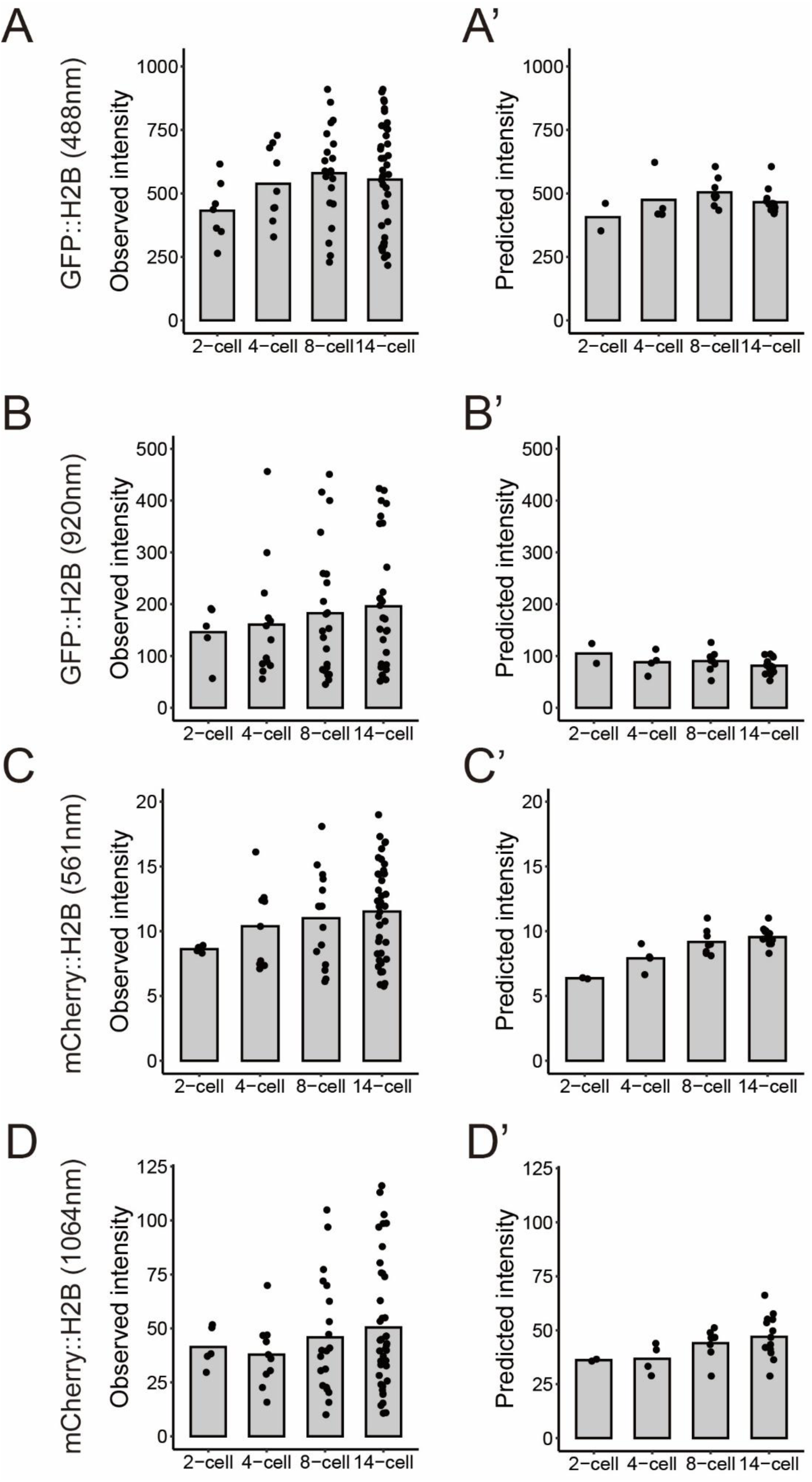
Developmental stage-dependent fluorescence intensity. (A-D) Observed fluorescence intensities at *T* = 0.5 (midpoint of the cell cycle) across developmental stages. (A) GFP::H2B imaged at 488 nm. (B) GFP::H2B imaged at 920 nm. (C) mCherry::H2B imaged at 561 nm. (D) mCherry::H2B imaged at 1064 nm. (A′–D′) Corrected fluorescence intensities including residual variation at *T* = 0.5 across developmental stages. Each point represents an individual cell, and bars indicate mean intensity for each stage. Distributions of fluorescence intensity within each stage and changes in mean intensity across stages can be compared between observed and corrected data.

Notably, after correction, the mean fluorescence intensity remained approximately constant across developmental stages (Fig. 7). This consistency was observed across all imaging conditions examined (GFP and mCherry, one-photon and two-photon), indicating that the result is robust to imaging modality and fluorophore.

This observation raises the possibility that histone levels are regulated to maintain a relatively constant concentration in the nucleus. In addition, the absence of systematic intensity decrease over time suggests that photobleaching was minimal under the imaging conditions used.

Together, these results indicate that removal of geometric bias enables more reliable estimation of underlying fluorescence levels, thereby increasing confidence in the observed convergence toward a constant intensity. This, in turn, may provide a basis for investigating mechanisms that regulate histone abundance, and suggests that the present framework could be broadly applicable to quantitative analysis of other fluorescently labeled molecules in developing systems.

## Discussion

### Microscopy insights and the need for computational correction

Fluorescence intensity in volumetric microscopy is strongly influenced by axial position (*z*), and potentially by radial position (*r*), making quantitative comparison across cells and samples inherently challenging. One potential strategy to mitigate this problem is to employ imaging modalities that reduce *z*-dependent signal attenuation. In this study, we therefore extended a spinning-disk confocal system to two-photon excitation [12–14] and, to our knowledge, provide one of the first demonstrations of its application to live imaging of embryonic development. Two-photon microscopy is widely regarded as advantageous for deep tissue imaging due to its reduced scattering and confinement of excitation to the focal volume [3,7]. Consistent with its optical properties, nuclei appeared less distorted along the *z*-axis, indicating improved optical sectioning, segmentation accuracy and more faithful representation of three-dimensional structure (Fig. 2E, 2F). However, these improvements did not eliminate *z*-dependent intensity bias, as fluorescence intensity still exhibited systematic attenuation along the *z*-axis (Fig. 2B, 2D). One contributing factor may be that spinning-disk systems rely on confocal detection through pinholes, which can reduce photon collection efficiency relative to non-descanned detection used in conventional two-photon systems, limiting the benefit of two-photon excitation for quantitative intensity measurements. These findings highlight that hardware improvements and computational correction should be regarded as complementary strategies, and that computational frameworks explicitly addressing geometric attenuation remain necessary when advanced imaging modalities are employed.

### Observed attenuation patterns are inconsistent with predictions from physical optical models

The parametric forms adopted in this study — polynomial terms in *z* and a linear term in *r* — were motivated by empirical model selection rather than physical first principles, and the results revealed informative deviations from simple optical expectations.

For axial attenuation, physical models based on Beer–Lambert-type absorption and scattering predict that fluorescence intensity decays exponentially with depth, which corresponds to a linear dependence of log-transformed intensity on z [11]. However, model selection consistently favored the inclusion of a quadratic z term across all datasets, indicating that the effective attenuation deviates from a simple exponential. This is consistent with the expectation that multiple depth-dependent processes — including changes in the point spread function, refractive index mismatch, and the interplay between excitation and emission scattering — accumulate in a way that cannot be captured by a single exponential decay. The adequacy of a low-order polynomial as an empirical approximation does not imply that these physical processes are absent; rather, it suggests that their combined effect can be approximated parsimoniously within the range of depths examined.

For radial attenuation, the expected behavior based on optical principles is also well defined. If the excitation beam has a Gaussian profile centered on the optical axis, illumination intensity falls off as exp(−*r*²). Under one-photon excitation, detected fluorescence is proportional to excitation intensity, so log-intensity would decrease quadratically with *r*. Under two-photon excitation, where fluorescence scales with the square of excitation intensity, the signal would be proportional to exp(−2*r*²), leading to the same qualitative prediction — a negative quadratic relationship between log-intensity and *r* — but with twice the magnitude. In either case, optical principles predict that intensity should decrease toward the periphery of the field of view. Instead, we observed a consistently positive linear *r* coefficient across all four datasets: apparent fluorescence intensity increased with radial distance from the optical axis. This result is qualitatively opposite in sign, ruling out simple illumination falloff as the primary determinant of radial variation in biological specimens.

The most likely explanation involves the geometry of the specimen itself. In our experimental configuration, the optical axis approximately coincides with the center of the embryo, where the specimen is thickest. Cells located near the center therefore experience greater scattering and absorption along the optical path than those at the periphery, where the path length through the sample is shorter. This reduction in scattering toward the periphery may compensate for, or even outweigh, the decrease in excitation intensity with radial distance, resulting in a net increase in apparent fluorescence. The spatial heterogeneity of optical properties within the embryo — including differences in refractive index between subcellular compartments (Goda et al., 2024) — may further complicate this relationship in ways not captured by simple physical models or calibration measurements on uniform samples. Notably, positive radial dependence was consistently observed across both one-photon and two-photon imaging conditions, despite their fundamentally different optical mechanisms, further supporting the view that the observed radial behavior reflects specimen-specific rather than system-level optical effects.

Together, these findings illustrate that neither the functional form nor the sign of position-dependent attenuation in biological specimens can be reliably predicted from optical models calibrated on homogeneous samples. This motivates the data-driven approach taken in this study, in which attenuation parameters are estimated directly from replicated observations of the biological specimen under realistic imaging conditions.

### Relationship to existing correction approaches

Our approach shares with methods for correcting in-plane illumination non-uniformity — including calibration-based flat-field correction using uniform fluorescent reference samples (Model and Burkhardt, 2001; Zwier et al., 2008) and retrospective methods such as BaSiC and CIDRE that infer correction fields from image collections without dedicated calibration images (Peng et al., 2017; Smith et al., 2015) — the general principle of estimating systematic, position-dependent imaging artifacts and removing them from the observed signal. However, these methods address spatial inhomogeneity within the focal plane (*xy*), rather than the axial (*z*) and radial (*r*) position-dependent attenuation that is the primary challenge in volumetric imaging of thick biological specimens. Furthermore, they rely on the assumption that fluorophore distributions are approximately uniform across images or reference samples, which may not hold in structured biological specimens. This distinction is critical: because attenuation parameters in our framework are inferred from biologically equivalent cells imaged at varying positions, the correction captures specimen-specific optical effects that system-level shading correction cannot account for — as directly demonstrated by the reversal of radial dependence between calibration samples and biological specimens.

Our approach is also distinct from empirical methods that estimate *z*-dependent decay from the mean intensity of each image plane [11], which assume fluorophore density to be uniform along the axial direction — an assumption rarely met in structured biological specimens — and from the calibration-based strategy [6], which requires a panel of reference strains and assumes constant expression levels across all cells. By contrast, our framework requires no external reference and infers attenuation directly from replicated observations of the specimen of interest, using the biological reproducibility of *C. elegans* embryogenesis as the internal reference. Model selection using AIC allows the functional form of attenuation — including nonlinear *z* dependence and the presence or absence of *r* dependence — to be determined empirically rather than assumed a priori.

### Biological implications

An unexpected observation of this study was that, after correction, mean histone fluorescence intensity remained approximately constant across developmental stages despite substantial changes in nuclear size. This behavior is not readily explained by simple expectations. If histone concentration reflects chromatin concentration, one would expect that, because the total amount of chromatin per nucleus remains approximately constant while nuclear volume decreases during development, histone concentration should increase. Conversely, if the total histone amount (*N*_tot_) is conserved (*C*_N_ *V*_N_ + *C*_C_ *V*_C_ = *N*_tot_, where *C*_N_, *C*_C_, *V*_N_ and *V*_C_ are the concentration (*C*) and volume (*V*) of nucleus (_N_) and cytoplasm (_C_), respectively) and nuclear–cytoplasmic exchange follows a simple equilibrium (*C*_N_ / *C*_C_ = *K* (constant) > 1), the increasing ratio of nuclear volume to cytoplasmic volume would be expected to reduce nuclear histone concentration (*C*_N_ = *K N*_tot_ / {(*K*-1) *V*_N_ + *V*_tot_}, where *V*_tot_ = *V*_N_ + *V*_C_). However, neither of these scenarios is consistent with the observed stage-invariant intensity. These considerations highlight that maintaining constant nuclear histone concentration is not trivially explained by simple physical or geometric models, suggesting that additional regulatory mechanisms may be involved.

More broadly, this finding illustrates an important implication of the present approach: by removing imaging-dependent biases, the correction framework reveals biological regularities that are otherwise obscured in raw fluorescence measurements. This capability is not limited to relatively uniform markers such as histones, but is broadly applicable to quantitative analysis of gene expression or protein localization in developing systems.

### Limitations and future directions

The present framework has two principal limitations that point toward directions for future development. First, the method relies on the assumption that cells sharing the same identity and cell-cycle stage carry equivalent intrinsic fluorescence, and on the ability to assign those identities unambiguously across replicate specimens. Both conditions are well satisfied in *C. elegans* embryos, where developmental reproducibility is exceptionally high and cell identities are defined by an invariant cell lineage. Extension to systems with greater cell-to-cell variability, stochastic developmental trajectories, or less stereotyped lineage relationships would require either a substantially larger number of replicate observations to average out biological variability, or the incorporation of additional covariates to account for known sources of heterogeneity. More fundamentally, identifying biologically equivalent cellular states across specimens — a prerequisite for applying the method — may itself require dedicated computational approaches in systems where a fixed cell lineage is not available.

Second, the current implementation operates at the level of segmented objects — in this study, individual nuclei — and estimates attenuation from object-level mean intensity measurements. This design is well suited to specimens where discrete, clearly segmentable structures can be defined, but is less directly applicable to continuous fluorescence distributions or specimens where cell boundaries are difficult to delineate. Adapting the framework to image-level or subcellular measurements would require modified formulations of the biological equivalence constraint.

Finally, while the present results demonstrate the utility of empirical, data-driven correction, a deeper integration with physical optical models could improve both the interpretability and the transferability of the estimated parameters across imaging systems. Physics-informed modeling frameworks, which embed mechanistic constraints within statistical estimation procedures, represent a promising direction for developing correction methods that combine the flexibility of data-driven approaches with the generalizability afforded by physical understanding.

## Materials and methods

### Worm culture and establishment of *C. elegans* strain

The worm strains used in this study are listed in Sup. Table S6. The worms were maintained under standard conditions [16]. CAL2641 was made by crossing CAL2501 (*unc-119*(*ed3*)III; *wjIs180* [*Pmex-5*::42B3::GFP::*Ttbb-2* + *unc-119*(+)]) and CAL2421 (*unc-119*(*ed3*)III; *wjIs108* [*Ppie-1*::mCherry::*his-58*::*pie-1* + *unc-119*(+)]) and selecting the F2 generation for having both homozygous for *wjIs180* and *wjIs108*. CAL2421 was made by back-crossing CAL0941 [17] with N2 and selecting the F2 generation for having both homozygous for *wjIs108*. CAL2501 contains the sequence of clone 42B3 (R78K/A80T/M95V) of RNAPII-S2P mintbody [18], which has been codon-optimized for nematodes. CAL0231 was previously reported [19].

### Sample preparation for imaging

Adult hermaphrodites were dissected in 0.75 × egg salt (118 mM NaCl, 40 mM KCl, 3.4 mM CaCl_2_, 3.4 mM MgCl_2_, 5 mM HEPES pH 7.2). A 30 µm-thick double-sided adhesive tape was applied to a glass slide such that the outer edge of the tape aligned with the perimeter of an 18 × 18 mm coverslip (Matsunami Glass Ind., Osaka, Japan). The embryo was placed on a poly-L-lysine (PLL) -coated coverslip, and 10 µL of 0.75 × egg salt was added. The coverslip was then inverted onto the slide and mounted. The edges of the chamber were sealed with VALAP (2:2:1 mixture of Vaseline, lanolin, and paraffin) to prevent evaporation and maintain a stable imaging environment.

### Microscopy setup and image acquisition

Live imaging was performed using a spinning-disk confocal microscope system (CSU-MP; Yokogawa Electric, Kanazawa, Japan) [13,14]. The system is based on the CSU-X1 platform and can be switched between single-photon excitation (488nm or 561nm) and two-photon excitation (920nm or 1064nm) with a manual lever. The CSU-MP scanner is equipped with 100-μm pinholes aligned with a Nipkow disk incorporating microlenses with a diameter of 580-μm, rotating at 5,000 rpm. The pinhole diameter and interpinhole distance were longer relative to the conventional CSU-X1 configuration [12]. This design reduces pinhole cross-talk—i.e., unintended transmission of out-of-focus fluorescence through neighboring pinholes—which is a major source of background noise when imaging thick specimens. Although this modification improves signal contrast in deep imaging, it also reduces confocal sectioning to some extent.

The scanner is mounted on an inverted microscope (IX71; Olympus, Tokyo, Japan) equipped with a 60x silicon-immersion objective lens (UPlanSApo, 60x/1.30Sil; Olympus) and relay lenses with a magnification of ×2.0. Z-stack imaging was performed using a piezo actuator (P-915K323; Physik Instrumente, Karlsruhe, Germany) controlled via a NIDAQ system (National Instruments, Austin, TX, USA). Z-stacks were acquired over a total thickness of 30.5 μm with a z-interval of 0.5 μm. Time-lapse imaging was performed at 2-minute intervals for 90 or 120 minutes, starting from the one-cell stage when the nucleus was positioned at the cell center. Images were acquired using an EM-CCD camera (iXon DU897-BV; Andor Technology, Belfast, UK), and the entire imaging system was controlled by NIS-Elements software (Nikon, Tokyo, Japan). The spatial resolution of the acquired image was 0.136 μm per pixel. Fluorescence imaging was performed using both one-photon and two-photon excitation under the following conditions. CAL0231 (GFP::H2B) was imaged using a 488 nm laser (Melles Griot, 85BCS-030-053, 30 mW) at 5% power with an exposure time of 200 ms for one-photon imaging, corresponding to approximately 1.6 mW measured before the objective lens. For two-photon imaging, a 920 nm laser (Spark lasers, ALCOR-920IR-2, averaged power: 2 W, pulse duration: 100 fs, repetition rate: 80 MHz) at 27% power with an exposure time of 400 ms, corresponding to approximately 150 mW before the objective lens. CAL2641 (mCherry::H2B) was imaged using a 561 nm laser (Cobolt, Jive, 25 mW) at 5% power with an exposure time of 60 ms for one-photon imaging, corresponding to approximately 0.04 mW before the objective lens. For two-photon imaging, a 1064 nm laser (Spark lasers, ALTAIR IR-5, averaged power: 5 W, pulse duration: 180 fs, repetition rate: 40 MHz) at 50% power with an exposure time of 96 ms, corresponding to approximately 700 mW before the objective lens. For two-photon excitation (920 nm and 1064 nm), the beam diameter (1.4mm) was expanded using a beam expander composed of a pair of plano-convex lenses (f = 25.4 mm and f = 125 mm; Thorlabs). These conditions were adjusted so that the nucleus could be perceived to the same degree. More than half of the imaged embryos successfully hatched, indicating that the imaging conditions did not substantially impair embryonic development.

### Nuclear segmentation

Nuclear regions were segmented from time-lapse 3D images of embryos using the surface detection function in Imaris (Bitplane). Surface detection was performed using the “background subtraction (local contrast)” option with a diameter of 2.5 µm. Nuclei were segmented based on the “Quality” index, with threshold values set to 110 for 488 nm images, 25 for 920 nm images, 3.0 for 561nm images, and 2.0 for 1064 nm images. In a small number of cases, minor adjustments to parameters were made to ensure accurate nuclear detection. Objects smaller than 500 voxels were excluded. Time-resolved tracking of nuclei was performed using the “lineage” tracking algorithm. The maximum displacement and maximum gap size were set to 4.08 µm and 2 frames, respectively.

Tracks were manually curated to ensure that each track corresponded to a single biological cell over time. Where necessary, segmentation and tracking errors were manually corrected using Imaris to maintain consistency of individual cell trajectories. After curation, each nucleus was assigned a cell identity based on lineage information. Surfaces that were not assigned a cell identity, including polar bodies, were excluded from subsequent analyses. In addition, if no corresponding surface was detected for a given cell identity at a specific time point, that observation was treated as missing and excluded from downstream analysis.

### Measurement of background intensity

Background intensity was measured using Fiji (ImageJ). For each image, the mean intensity was calculated from four square regions (40 × 40 pixels) located at the corners of the field of view, which were assumed to be outside the embryo. The average of these measurements was used as background intensity and subtracted from the nuclear fluorescence signal. No substantial differences in background intensity were observed across imaging wavelengths.

### Data preprocessing

All data preprocessing was performed using custom scripts in R (R Foundation for Statistical Computing). Measurements from each embryo were compiled into a unified dataset, with individual nuclei tracked over time. The radial position *r* was defined as the distance from the center of the image, which coincides with both the optical axis and the center of the embryo.

For each cell, time points (*f*) were defined based on frame indices obtained from tracking. To enable comparison across cells, time was normalized relative to the cell cycle. For a given cell, let *f_0_* denote the final frame index of its parent cell, and let *N* denote the total number of frames in the cell cycle of the cell. The cell was then tracked from frame *f* = *f_0_* + 1 to *f* = *f_0_* + *N*, and a normalized time variable *T* was defined. Under this definition, the first observed frame corresponds to *T* = 1 / *N*, and the last frame prior to nuclear division (i.e. anaphase onset) corresponds to *T* = 1. This normalization allows comparison of temporal dynamics across cells with different cell cycle lengths.

Data from multiple embryos were processed in a consistent manner and combined into a single dataset for downstream analysis. The resulting dataset consisted of time-resolved, single-nucleus measurements including mean fluorescence intensity within each segmented nucleus, the centroid coordinates of each nucleus, time, and cell identity. Specifically, for each observation *i*, we defined the measured mean fluorescence intensity *I*_i_, radial position *r*_i_ (i.e., distance from the center of the field of view in the x-y plane), axial position *z*_i_ (i.e., imaging depth along the optical axis), normalized time *T*_i_, and cell identity *c*(*i*), which were used for downstream analysis.

### Statistical model for correcting intensity attenuation

We assumed that the observed fluorescent intensity is not solely determined by the underlying molecular abundance but is also systematically influenced by the spatial position of the cell within the embryo and its progression through the cell cycle. To account for these effects, we modeled the observed intensity *I*_i_ as a function of axial position (*z*), radial position (*r*) and cell cycle progression (*T*). The full model was defined as:

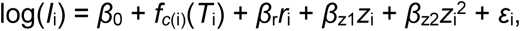

where *f_c_*_(i)_(*T*_i_) represents a cell-specific temporal effect modeled using natural cubic splines, and *ε*_i_ represents residual variation. We evaluated a set of candidate models by considering all combinations obtained by excluding the *r*_i_ term, the quadratic *z*_i_ term (*z*_i_^2^), or both *z*_i_ terms (*z*_i_ and *z*_i_^2^). For the spline term, we considered models with two or three internal knots. This class of formulations allows separation of geometric attenuation effects from biological variation using observational data. Model parameters were estimated by linear regression on log-transformed fluorescence intensity using the *lm* function implemented in R.

### Model selection

To determine the appropriate model structure, we performed systematic model selection by evaluating candidate models with different combinations of radial, axial, and temporal terms. Axial dependence was modeled as (i) no dependence, (ii) linear dependence, or (iii) quadratic dependence (*z* and *z*^2^). Radial dependence was modeled as either (i) no dependence or (ii) linear dependence. Temporal effects were modeled using natural cubic splines with (i) two or (ii) three internal knots. In total, 12 candidate models representing all combinations of these components were evaluated, and the best-fitting model was selected based on the Akaike Information Criterion (AIC).

### Intensity correction and prediction

Corrected fluorescence intensities were obtained using the fitted model. The corrected intensity, removing measurement noise and geometric attenuation effects, at the defined position (e.g. *r*_0_ = 0, *z*_0_ equal to the mean axial position within the embryo) for each observation was defined as:

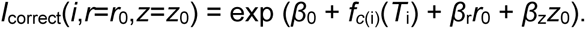

Residual analysis was used to assess whether systematic dependencies on radial (*r*) and axial (*z*) positions remained after correction.

## Acknowledgements

We are grateful to Tomoko Ozawa for assistance with nuclear segmentation; Hiroshi Nakayama (Yokogawa Electric) for designing CSU-MP system used in this study; Takashi Murata (Kanagawa Inst Tech) and Kentaro Kobayashi (Hokkaido Univ) for their advice and support on the CSU-MP system; Hiroshi Kimura (Science Tokyo) for kindly providing the plasmid sequence of the mintbody; Osamu Yasuhiko (Hamamatsu Photonics) for helpful discussion. This work was supported by the Data-Scientist-Type Researcher Training Project of The Graduate University for Advanced Studies, SOKENDAI for SI, by “Strategic Research Projects” grant from ROIS (Research Organization of Information and Systems) for AK, SI, SK and SA, by JSPS KAKENHI Grant Numbers JP16H06280 (“Advanced Bioimaging Support”), 18H05529, 23H04294, 25K02276 for AK, 25K15034 for SK, and 25H01025 for TN and KO, by Japan Agency for Medical Research and Development (AMED) Brain/MINDS 2.0, JP24wm0625105 for TN and KO, and Japan Science and Technology Agency (JST) FOREST JPMJFR230R for KO.

## Author contributions

S.I. and A.K. conceived the project. S.I. developed the model under the supervision of S.K. and S.A. K.O. and T.N. contributed to the implementation of the two-photon spinning-disk microscopy system. S.I. performed all experiments and data analyses. S.I. and A.K. wrote the manuscript with input from all authors.

