## Supplemental Information for "A Data-Driven Correction Framework for Axial- and Radial-Position-Dependent Intensity Attenuation in Volumetric Fluorescence Microscopy"

### Supporting Information

**Supplementary Tables S1-S4: Model comparison across candidate attenuation models for each imaging condition.** Model comparison across 12 candidate models fitted to fluorescence intensity data for each imaging condition. Models were evaluated using Akaike Information Criterion (AIC), and are listed in ascending order of AIC for each dataset. Coefficient of determination ($R^2$), and adjusted $R^2$ are also shown. Each table corresponds to a different imaging condition (GFP::H2B at 488 nm (S1) and 920 nm (S2); mCherry::H2B at 561 nm (S3) and 1064 nm (S4)). The candidate models differ in their treatment of depth dependence (z_mode: no dependence (z0), linear (z1), or quadratic(z12)), radial dependence (r_mode: no dependence (r0), linear (r1)), and temporal dynamics (number of spline knots, n_internal_knots). The number of parameters (k), sample size (n), and ΔAIC relative to the best model within each dataset are also reported. These comparisons were used to select the best-fitting model for subsequent analyses.

**Supplementary Table S5: Partial $R^2$ values for depth and radial terms across imaging conditions.** Partial $R^2$ values were calculated to quantify the contribution of depth ($z$) and radial position ($r$) to fluorescence intensity variation in the best-fitting model for each dataset (GFP::H2B at 488 nm and 920 nm; mCherry::H2B at 561 nm and 1064 nm). Partial $R^2$ was computed based on differences in $R^2$ between nested models. Across imaging conditions, depth ($z$) consistently explained a substantial proportion of variance, whereas radial position ($r$) contributed to smaller effects. In the 1064 nm dataset, the partial $R^2$ for the radial term was close to zero (slightly negative), indicating that its contribution was minimal and near the limit of detectability, although model selection criteria (AIC) supported its inclusion.

**Supplementary Table S6: Strain list.**

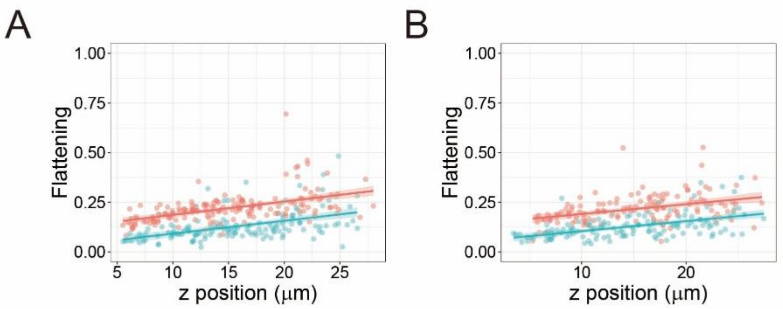


**Supplementary Figure S1: Nuclear flattening as a function of axial position (*z*).**

Scatter plots of nuclear flattening versus z-position for individual nuclei at the 14-cell stage ($T$ = 0.5). These data correspond to the unbinned measurements underlying Fig. 2E and 2F. (A) GFP::H2B imaged at 488 nm (one-photon, pink) and 920 nm (two-photon, light blue). (B) mCherry::H2B imaged at 561 nm (one-photon, pink) and 1064 nm (two-photon, light blue). Lines indicate linear regression fits.

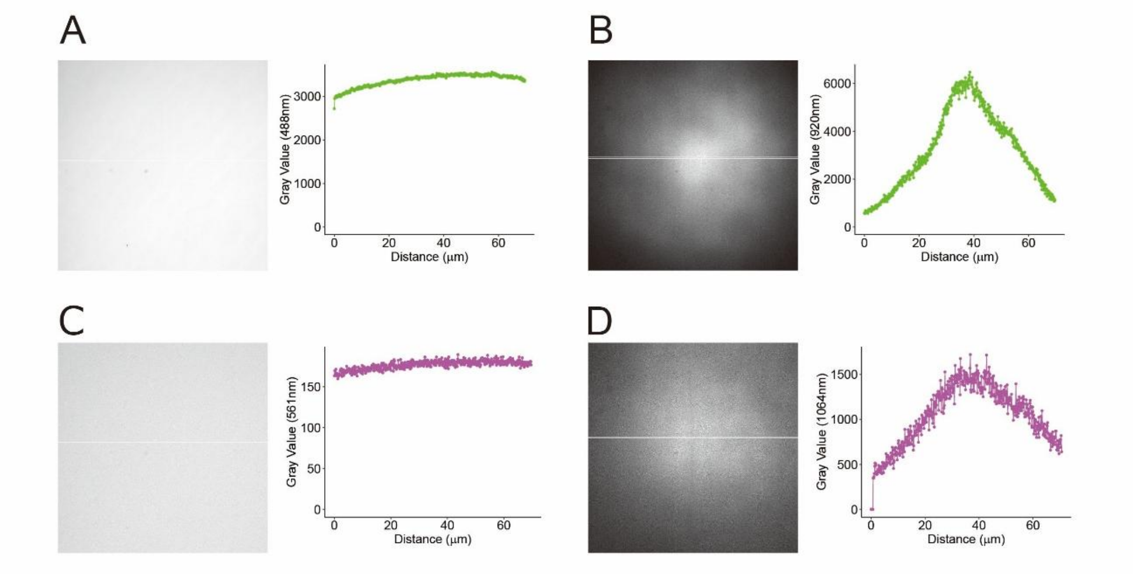


**Supplementary Figure S2: Radial dependence of fluorescence intensity in a homogeneous fluorescent sample.**

Representative images of a homogeneous fluorescent plate and corresponding intensity profiles across the field of view. For each panel, the left image shows the fluorescence intensity in grayscale, and the right plot shows the intensity profile measured along a horizontal line passing through the center of the image.

(A) 488 nm excitation. (B) 920 nm excitation. (C) 561 nm excitation. (D) 1064 nm excitation. In all cases, fluorescence intensity decreases toward the periphery of the field of view, indicating radial attenuation even within a single focal plane.

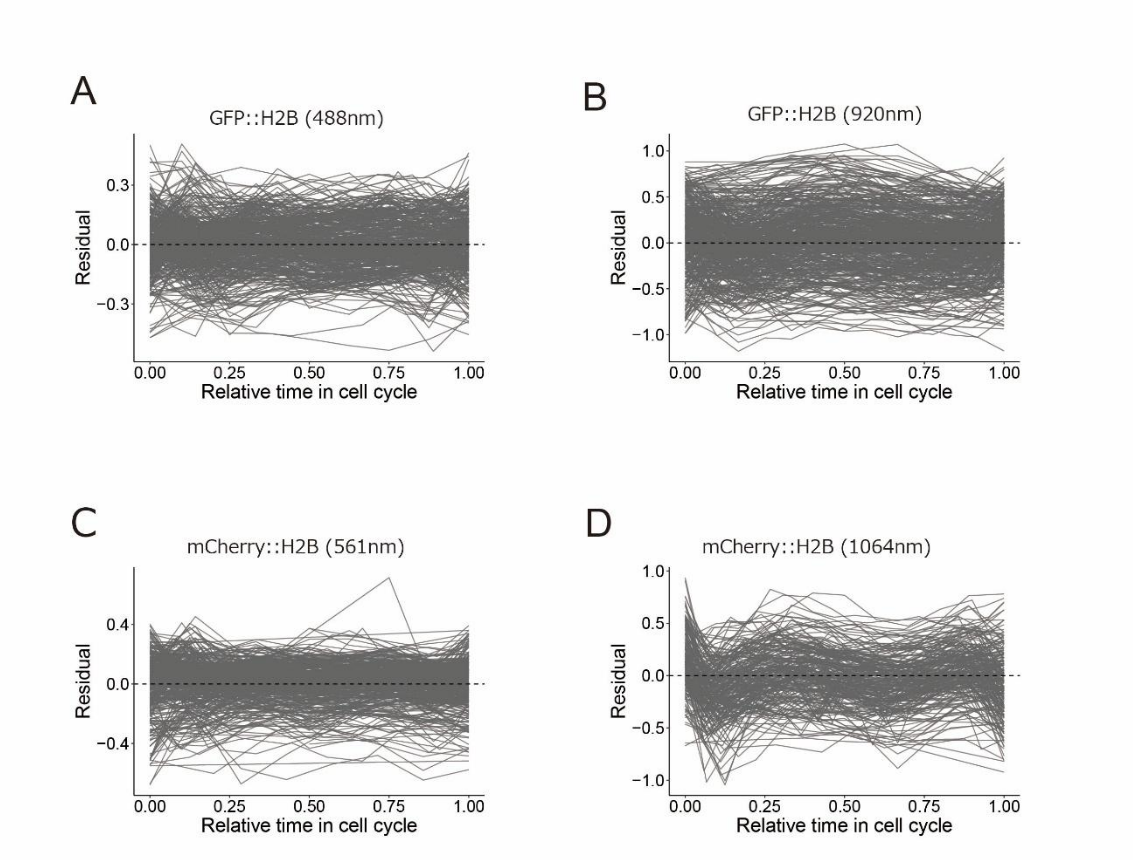


**Supplementary Figure S3: Residual dependence on normalized cell cycle time.** Residuals from the best-fitting model (as shown in Fig. 6) plotted against normalized time (*T*), showing no systematic temporal trends. Each line represents a cell-derived datapoint. (A) GFP::H2B imaged at 488 nm. (B) GFP::H2B imaged at 920 nm. (C) mCherry::H2B imaged at 561 nm. (D) mCherry::H2B imaged at 1064 nm.

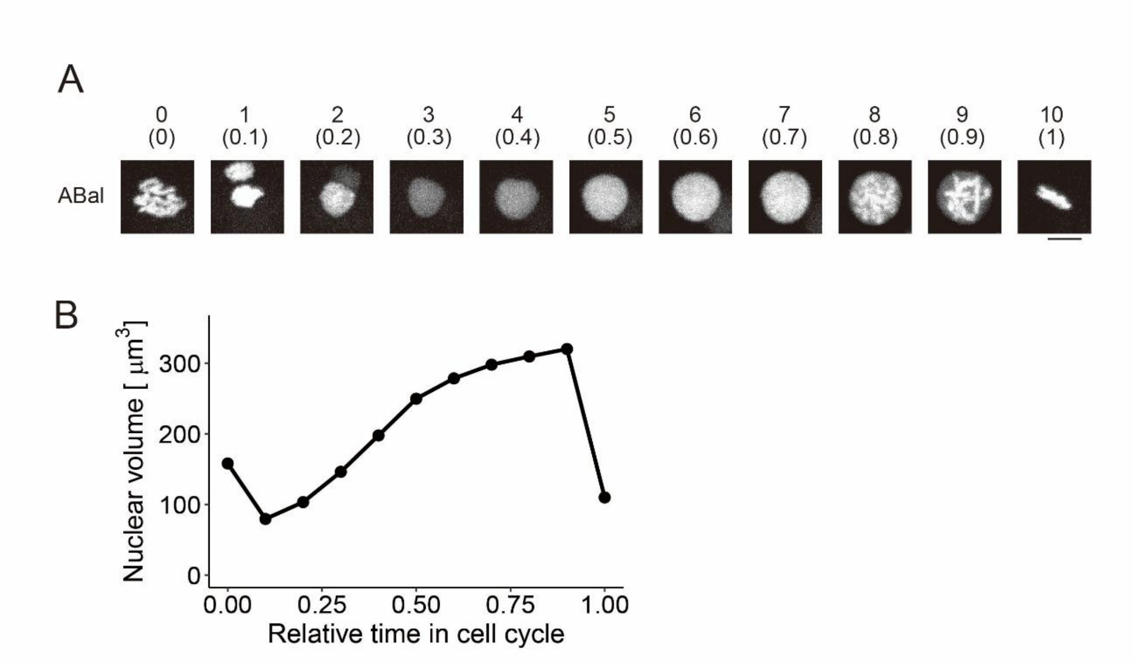


**Supplementary Figure S4: Histone and nuclear dynamics during cell cycle**

(A) Representative images of H2B. Histone fluorescence in the ABal cell at the eight-cell stage is shown from one anaphase to the next. Images were acquired at 2-minute intervals. Numbers indicate frame indices within the cell cycle, with normalized time shown in parentheses. Scale bar: 5 μm. (B) Nuclear volume dynamics of the ABal cell. Data in (A, B) were obtained using GFP::H2B imaged at 920 nm.
